# Towards reproducible models of sequence learning: replication and analysis of a modular spiking network with reward-based learning

**DOI:** 10.1101/2023.01.18.524604

**Authors:** Barna Zajzon, Renato Duarte, Abigail Morrison

## Abstract

To acquire statistical regularities from the world, the brain must reliably process, and learn from, spatiotemporally structured information. Although an increasing number of computational models have attempted to explain how such sequence learning may be implemented in the neural hardware, many remain limited in functionality or lack biophysical plausibility. If we are to harvest the knowledge within these models and arrive at a deeper mechanistic understanding of sequential processing in cortical circuits, it is critical that the models and their findings are accessible, reproducible, and quantitatively comparable. Here we illustrate the importance of these aspects by providing a thorough investigation of a recent model proposed by Cone and Shouval (2021). We re-implement the modular columnar architecture and reward-based learning rule in the open-source NEST simulator, and successfully replicate the main findings of the original study. Building on these, we perform an in-depth analysis of the model’s robustness to parameter settings and underlying assumptions, highlighting its strengths and weaknesses. We demonstrate a limitation of the model consisting in the hard-wiring of the sequence order in the connectivity patterns, and suggest possible solutions. Finally, we show that the core functionality of the model is retained under more biologically-plausible constraints.

## 1 Introduction

Navigating in a dynamic environment requires actions and decisions that are precisely coordinated in time and space, matching the spatio-temporally structured stimuli upon which they are based. Therefore, the ability to learn, process and predict sequential patterns is a critical component of cognition, with recent experimental findings showing a multitude of brain regions to be involved in sequence processing (Dehaene et al., 2015; Wilson et al., 2018; Henin et al., 2021). Some areas, such as the hippocampus, specialize on (spatial) tasks that rely mainly on the ordinal information within the sequence and *compress* the temporal features, for instance by recalling sequences faster than experienced (August and Levy, 1999). Other regions, including early sensory areas such as the primary visual cortex, are capable of learning and recalling not just the order of a series of stimulus patterns, but also the duration of the individual elements (Xu et al., 2012; Gavornik and Bear, 2014). In fact, the ability to represent both the ordinal and temporal components of a sequence are two of the most fundamental requirements for any system processing sequential information.

However, most existing models of unsupervised biological sequence learning address only the first of these two criteria, focusing on acquiring the order of elements and typically failing to account for their duration. They either cannot intrinsically represent the time intervals (Klos et al., 2018; Bouhadjar et al., 2021), or they assume a fixed and identical duration for each element that is limited by the architecture (Maes et al., 2021), or they produce longer sequences that arise spontaneously even in the absence of structured input (and hence are not related to it, Fiete et al., 2010). Other studies have shown that event and stimulus duration can be encoded via transient trajectories in the neural space through the sequential activation of different cell assemblies, but these mechanisms were either restricted in time (Duarte and Morrison, 2014; Duarte et al., 2018), explored in the context of working memory (Mongillo et al., 2008; Fitz et al., 2020) or relied on heavily engineered network architectures (Klampfl and Maass, 2013).

Seeking to unify these computational features, Cone and Shouval (2021) recently proposed a novel, biophysically realistic spiking network model that avoids the problem of temporal compression while maintaining the precise order of elements during sequence replay. Relying on a laminar structure, as well as experimentally observed cell properties, the system uses a local, eligibility-based plasticity rule (3-factor learning rule see, e.g. Frémaux et al., 2015; Porr and Wörgötter, 2007; Magee and Grienberger, 2020; Gerstner et al., 2018) to learn the order of elements by mapping out a physical path between stimulus-tuned columns (akin to (Zajzon et al., 2019)), with the duration of each item being encoded in the recurrent activations within the corresponding column. The learning rule, based on the competition between two eligibility traces and a globally available reward signal, is grounded in recent experimental findings (He et al., 2015; Huertas et al., 2016). This modular architecture allows the network to flexibly learn and recall sequences of up to eight elements with variable length, but only with simple transitions between items (first-order Markovian). More intricate sequences with history dependence (i.e., higher-order Markovian) can be learned, but require additional structures for memory. Given the increased complexity, this ability is only demonstrated in a continuous rate-based model.

The code for the model is available in *MATLAB*. As this is a proprietary, closed-source software, models expressed in this manner have accessibility issues (not every scientist can afford a license) and bear a greater risk of becoming non-executable legacy code, if the code is not regularly maintained (for an example, see Schulte to Brinke et al, 2022). Additionally, as *MATLAB* is a general purpose numeric computing platform, the researcher must develop all neuroscientific models and simulation algorithms *de novo*, which presents a higher risk for implementation errors and poorly-suited numerics (Pauli et al., 2018).

In this article we therefore present a replication of the original study, which serves the twin purpose of testing the original findings and providing a more accessible version of the model to the computational neuroscience community. Specifically, we re-implement their model using the open source software *NEST* (Gewaltig and Diesmann, 2007) to simulate the networks and Python for data analysis, thus ensuring a reusable and maintainable code base.

Here, we use the term *replication* in the R^5^ sense described by Benureau and Rougier (2018), i.e. striving to obtain the same results using an independent code base, whereas a *reproduction* (R^3^) of the model would have been achieved if we had obtained the results of the original study using the original code. However, others have argued these terms should be used the other way around: see Plesser (2018) for an overview and analysis.

Our re-implementation successfully replicates the principal results on the spiking network model from the original publication. Going beyond the reported findings, we perform an extensive sensitivity analysis of the network and learning parameters, and identify the critical components and assumptions of the model. We test the model at multiple scales and infer basic relations between the scale and numerical values of different parameters.

Additionally, we show that the original model and implementation rely on pre-wired feedforward projections between the columns to successfully learn the order of elements within a given sequence. We discuss why learning fails when generalizing to a more plausible architecture in which projections between all columns are allowed, and provide two possible solutions which restore the system’s functionality. Finally, we demonstrate that the core learning mechanisms can be retained in a functionally equivalent network architecture that contains only local inhibitory circuits, in line with cortical connectivity patterns (Brown and Hestrin, 2009).

The challenges we faced in replicating this study highlight the importance of detailed and accurate documentation, as well as access to the model code. In fact, a successful replication of the main results would not have been possible without being able to refer to the original implementation. In addition to multiple discrepancies between the model description and the code, some of the conceptual limitations we reveal here arise from certain critical implementation details (as discussed in Pauli et al., 2018).

Our findings thus demonstrate that undertakings such as these to replicate a study can also serve to improve the overall quality and rigour of scientific work. Moreover, if carried out shortly after the original publication, such in-depth analysis can lead to a better understanding of the computational model and thus both increase the likelihood that further models will be based on it, and decrease the likelihood that those models contain incorrect implementations or implicit (but critical) assumptions.

## 2 Results

To investigate how temporal sequences of variable durations can be acquired by cortical circuits, Cone and Shouval (2021) propose a chain-like modular architecture where each population (module) is tuned to a specific element in the sequence, and learning translates to modifications of the synaptic weights within and between modules, based on reward signals. We re-implement the model, originally in *MATLAB*, using the open-source software *NEST*. For access to the original code and our re-implemented version, please see the Data Availability Statement below.

The model is schematically illustrated in Figure 1A). Following a training period where the modules are stimulated in a particular order over multiple trials, the network should be able to recall/replay the complete sequence from a single cue. If learning was successful, both the order and duration of the elements can be recalled faithfully.

**Figure 1:**
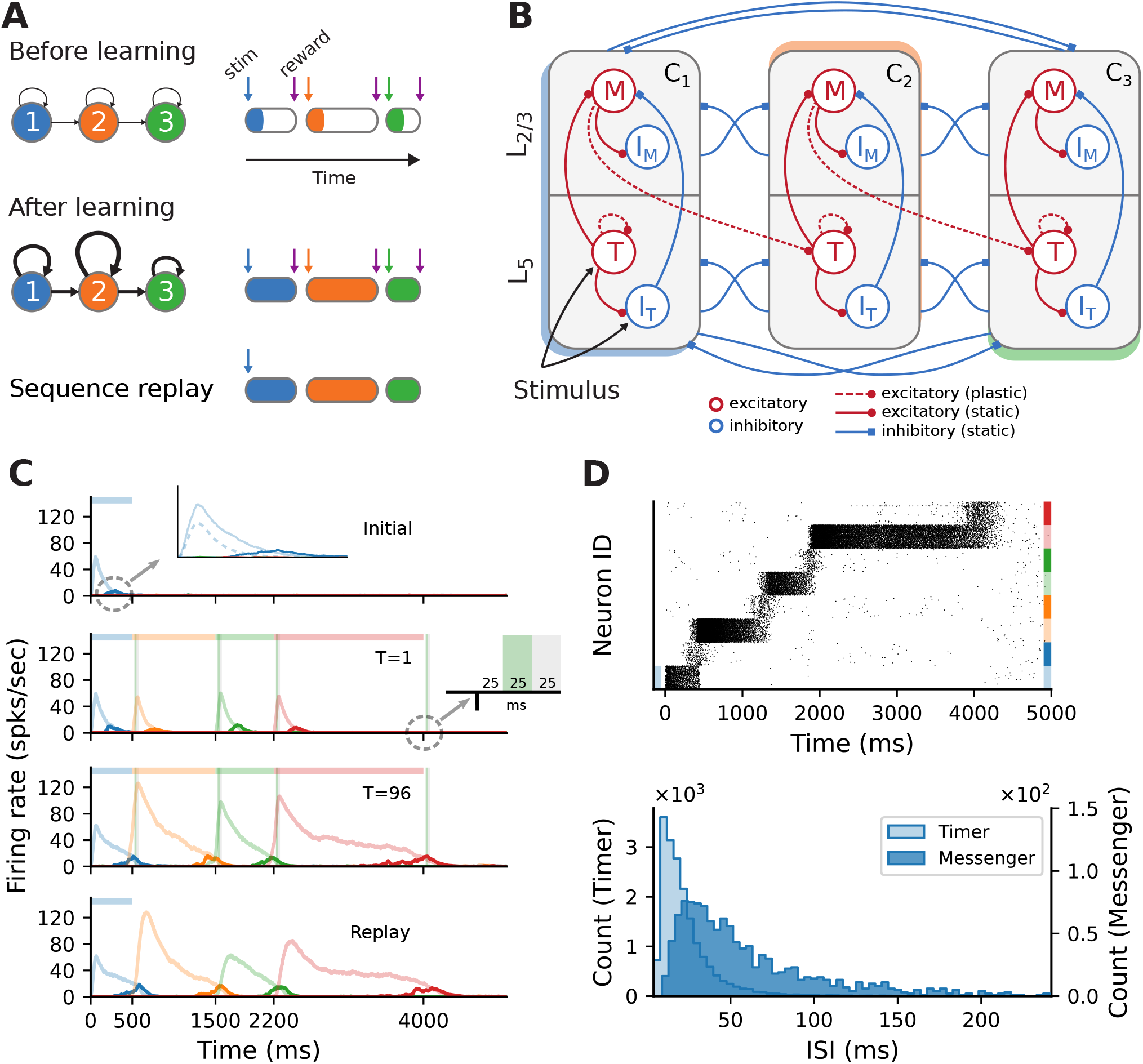
Sequence learning task and network architecture. **(A)** A sequence of three intervals (elements) is learned by a network with as many dedicated populations (columns). The individual populations are stimulated sequentially, with a global reward signal given at the beginning and the end of each element. After training, the recurrent and feedforward weights are strengthened, and the sequence is successfully recalled following a cue. The fullness of the colored sections on the right illustrates the duration of the activity (firing rates) above a certain threshold. **(B)** Each stimulus-specific column is composed of two excitatory, *Timers* (*T*) and *Messengers*(*M*), and two corresponding inhibitory populations, *I*_T_ and *I*_M_. Solid (dashed) arrows represent fixed static (plastic) connections. Cross-columnar inhibition always targets the excitatory population in the corresponding layer (*L*_5_ or *L*_2*/*3_). **(C)** Firing rates of the excitatory populations during learning (top three plots) and recall (bottom plot) of four time intervals (500, 1000, 700, 1800 ms). Light (dark) colors represent *T* (*M*) cells. Dashed light blue curve in top panel inset shows the inhibitory population *I*_T_ in *L*_5_. Green (grey) vertical bars show the 25 ms reward (trace refractory) period, 25 ms after stimulus offset (see inset). **(D)** Spiking activity of excitatory cells (top) and corresponding ISI distributions (bottom), during recall, for the network in **(C)**. In the raster plot, neurons are sorted by population (*T, M*) and sequentially by column (see color coding on the right).

Initially, each module exhibits only a transient activity in response to a brief stimulus (50 ms, see Methods), as the connections are relatively weak. The duration of each sequence element is marked by a globally available reward signal, forming the central component of a local reinforcement learning rule based on two competing, Hebbian-modulated eligibility traces (Huertas et al., 2016). This synapse-specific rule is used to update the weights of both recurrent and feedforward connections, responsible for the duration of and transition between elements, respectively. After learning, these weights are differentially strengthened, such that during a cued recall the recurrent activity encodes the current element’s extent, while the feedforward projections stimulate the module associated with the next sequence element.

The modules correspond to a simplified columnar structure roughly mapping to L2/3 and L5 in the cortex. The columns are composed of two excitatory populations, *Timer* (*T*) and *Messenger* (*M*), and two associated inhibitory populations *I*_T_ and *I*_M_ (Figure 1B), each containing 100 LIF neurons and conductance-based, saturating synapses (see Methods). Timer cells learn to represent the duration through plastic recurrent connections, while Messenger cells learn the transitions to the column associated with the next sequence element. Note that, unless otherwise mentioned, feedforward projections exist only between columns corresponding to consecutive items in the input sequence. In other words, the sequence transitions are physically traced out from the onset, only the weights are learned (see also Discussion). Cross-inhibition between the columns gives rise to a soft winner-take-all (WTA) behavior, ensuring that only one column dominates the activity.

### 2.1 Sequence learning and recall

This modular architecture allows the system to robustly learn and recall input sequences with variable temporal spans. Figure 1C depicts the population responses before and after the network has learned four time intervals, 500, 1000, 700 and 1800 ms (see also Figure 3 in Cone and Shouval, 2021). At first, stimulation of one column produces a brief response, with initial transients in the stimulated Timer and *L*_5_ inhibitory cells *I*_T_ (see Figure 1C, top panel and inset). With the inhibitory firing rate decaying faster than the Timers’ due to higher threshold and lack of recurrence (see Methods), there is a short window when the net excitation from the Timer cells elicit stronger responses from the Messenger cells.

**Figure 2:**
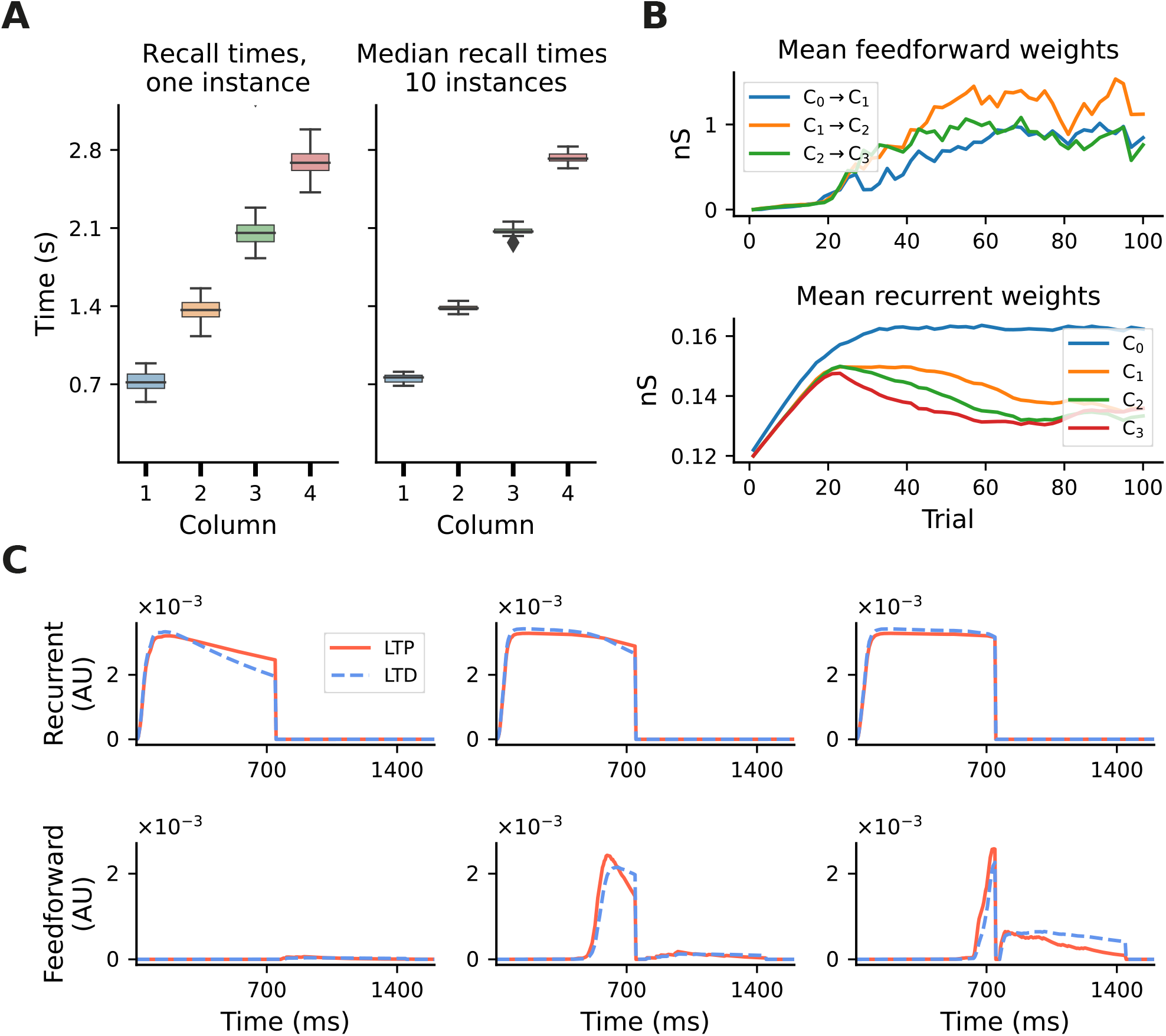
Accuracy of recall and evolution of learning. Results shown for a sequence of four intervals of 700 ms. **(A)** Fluctuations in learning and sequence recall. We define *recall time* as the time at which the rate of the Timer population drops below 10 spks*/*s. Left: recall times for 30 trials after learning, for one network instance. Right: distribution of the median recall times over 10 network instances, with the median in each network calculated over 30 replay trials. **(B)** Mean synaptic weights for feedforward (Messenger to Timer in subsequent columns, top) and recurrent (Timer to Timer in the same column, bottom) connections for onenetwork instance. **(C)** Mean LTP and LTD traces for the recurrent (top) and feedforward (bottom) connections, for learning trials T= 3, T= 15 and T= 35 and one network instance.

**Figure 3:**
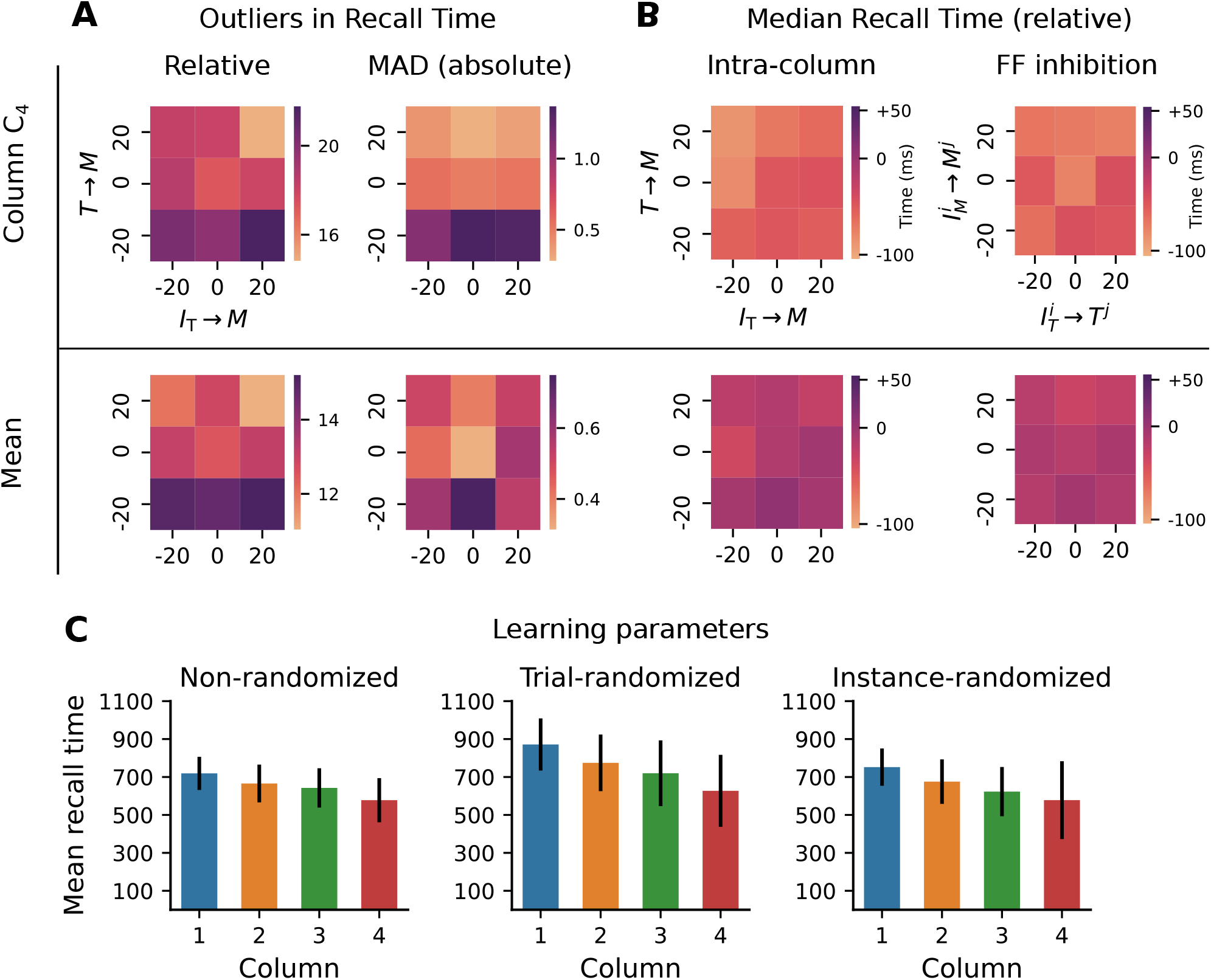
Robustness to variation in synaptic weights and learning parameters. The system was trained on a sequence of four elements, each with a duration of 700 ms. For the Timer cells, we define *relative recall time* as the recall time relative to stimulation onset, i.e., the time from the expected onset time (0, 700, 1400, 2100) in the sequence until the rate drops below a threshold of 10 spks*/*s. Conversely, *absolute recall time* is simply the time when the rate drops below threshold (relative to 0). **(A)** Number of outlier intervals reported during 50 recall trials, as a function of the percentage change of two synaptic weights within a column: excitatory Timer to Messenger, and inhibitory *I*_T_to Messenger. Top row shows the number of outliers, defined as a deviation of *±* 140 ms from the correct interval relative to expected onset (left), and the number of outliers detected using a modified z-score (threshold *>* 3, right panel) based on the median absolute deviation in column *C*_4_ (see main text). Bottom row shows the respective outliers averaged over all four columns. **(B)** Deviation of the median recall time from the expected 700 ms, as a function of the excitatory and inhibitory synaptic weights onto theMessenger cells in a column (left), and as a function of the cross-columnar (*C*_*i*_ ≀ *C*_*j*_) inhibitory synaptic weights within the same layers (right). Top and bottom row as in **(A)**. All data in **(A)** and **(B)** is averaged over 20 network instances. **(C)** Mean recall time of a four-element sequence of 700 ms intervals, over 50 recall trials of a single network instance. Left: baseline network. Center: during each training trial, the learning parameters (see main text) are drawn randomly and independently from a distribution of *±* 20% around their baseline value. Error bars represent the standard deviation. Right: the set of learning parameters is drawn randomly once for each network instance, with data shown averaged over 10 instances.

During training, when each column is stimulated sequentially, the recurrent Timer projections are strengthened such that their responses extend up to the respective reward signal (green vertical bars). At the same time, the feedforward projections from the Messenger cells on to the next column are also enhanced, such that upon recall (stimulation of first column), they are sufficient to trigger a strong response in the corresponding Timer cells.

This chain reaction allows a complete replay of the original sequence, preserving both the order and intervals. The activity propagation during recall is illustrated in Figure 1D (see Figure 3S4 in Cone and Shouval, 2021). The network displays realistic spiking statistics (coefficient of variation of 1.35 and 0.95 for Timer and Messenger cells), with Messenger cells having lower firing rates than Timer cells, roughly consistent with the experimentally observed values (Liu et al., 2015).

### 2.2 Learning and recall precision

The model exhibits fluctuations in the learning process and recall accuracy of sequences as a consequence of noise and the stochastic nature of spiking networks. For sequences of intermediate length, the recall times typically vary within *±* 10-15% of the target duration (see Figure 2A, left). However, this range depends on several parameters, and generally increases with duration or sequence length (see Supplementary Figure S1). Nevertheless, averaged over multiple network instances, these effects are attenuated and learning becomes more precise (Figure 2A, right).

These fluctuations can also be observed at the level of synaptic weights. Whereas the recurrent weights in the Timer populations converge to a relatively stable value after about 70 trials (see Figure 2B, bottom panel, and Figure 3S2 in Cone and Shouval, 2021), the feedforward weights display a larger variability throughout training (top panel). For the recurrent connections, convergence to a fixed point in learning can be formally demonstrated (see proof in Cone and Shouval, 2021). As a Hebbian learning rule (see Methods), the two competing LTP and LTD eligibility traces are activated upon recurrent activity in the Timer population. Assuming that both traces saturate quickly, with a slightly higher LTD peak, and given a larger time constant for the LTP trace, the LTD trace will decay sooner, resulting in the facilitation of recurrent synapses during the reward period (Figure 2C, top panel). Learning converges when the net difference between the two traces is zero at the time of reward.

For the feedforward weights, an analytical solution is more difficult to derive. Due to Hebbian co-activation of Messenger cells and Timer cells in the subsequent module, the traces are activated (non-zero) shortly before the reward period, temporarily reset following reward, and reactivated during the next trial (Figure 2C, bottom panel). The net weight change is thus the sum of trace differences over two subsequent reward periods. Empirically, learning nevertheless tends to converge to some relatively stable value if feedforward projections only exist between columns coding for subsequent input elements. However, because the reward signal is globally available at each synapse, all projections from a Messenger population to any other module could, in theory, be facilitated, as long as there is some temporal co-activation. We elaborate on this aspect in Section 2.5.

### 2.3 Model robustness

Although formally learning convergence is only guaranteed for the recurrent Timer connections, Cone and Shouval (2021) report that in practice the model behaves robustly to variation of some connectivity and learning parameters. However, the range of parameter values and sequence lengths analyzed in Cone and Shouval (2021) (see their Figure 5 and supplements) does not give a complete account of the parameters’ influence and the model’s limits. To test model robustness more thoroughly, we varied a number of the synaptic weights and learning parameters beyond those considered in the original work, and measured the consistency in the recall times of a sequence composed of four 700 ms intervals.

**Figure 4:**
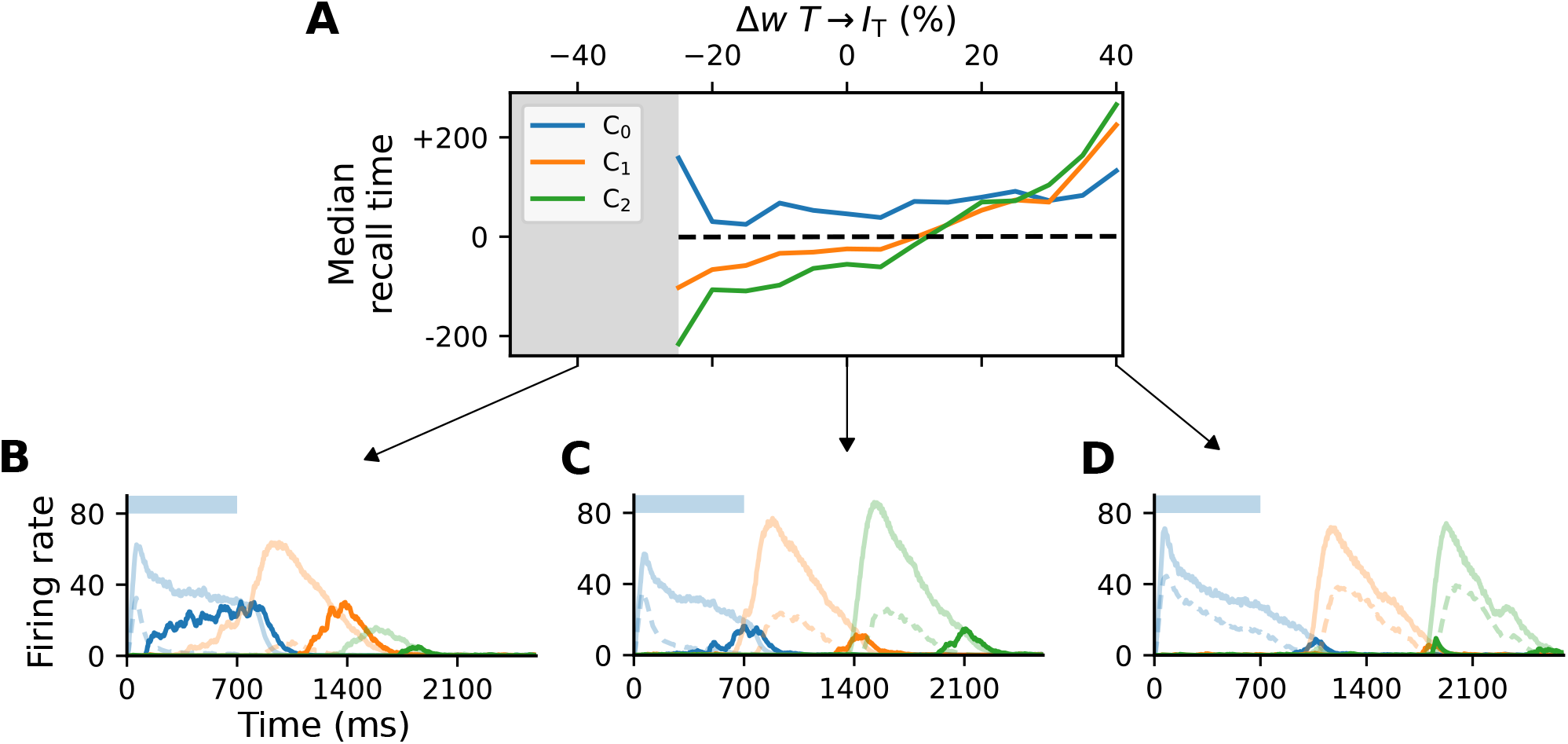
Activity of L_5_ inhibitory population is critical for accurate learning. **(A)** Deviation of the median recall time of three intervals of 700 ms, as a function of the change in synaptic weights *T→ I*_T_ relative to baseline (∆*w* = 0). Grey area (*<−* 25%) marks region where learning is unstable (not all elements can be recalled robustly). Data is averaged over 5 network instances. **(B-D)** Characteristic firing rates during recall for values deviations of *−* 25, 0 and 40% relative to baseline. Solid curves represent the excitatory populations as in Figure 1, while dashed curves indicate the respective inhibitory populations *I*_T_in *C*_*i*_.

**Figure 5:**
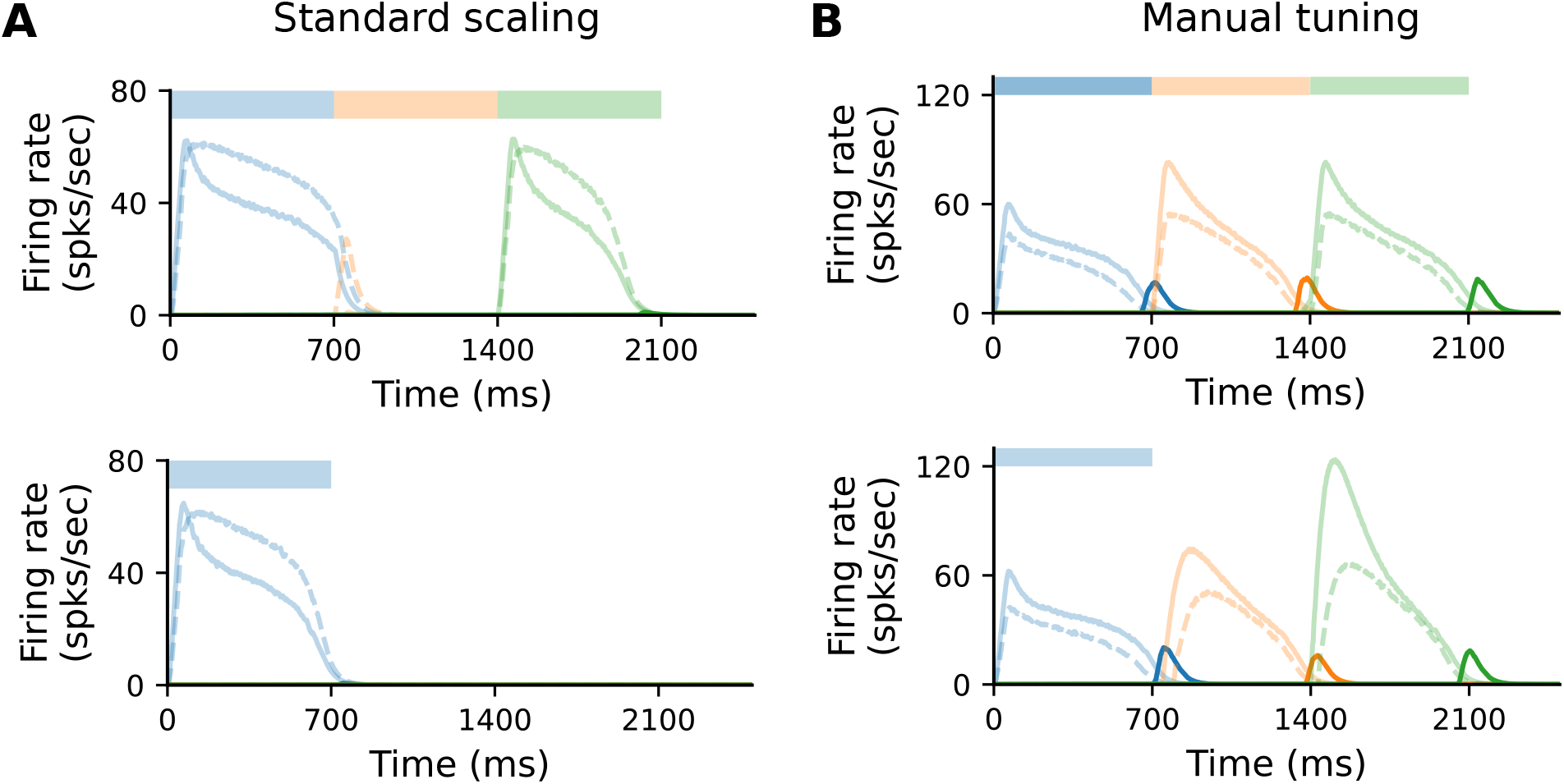
Scaling the model requires manual retuning of parameters. **(A)** Characteristic firing rates during training (top) and recall (bottom) of a sequence composed of three 700 ms intervals, in a larger network where each population is composed of *N*^*′*^ = 400 cells. All static weights have been scaled down by 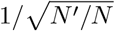 (see Methods). Solid curves show Timer (light) and Messenger (dark) cells, dashed curves *I*_T_cells. **(B)** As in (A), with further manual tuning of specific weights. For details, see Methods and Supplementary Material.

First, we varied the excitatory and inhibitory projections onto Messenger cells within a column, in an interval of *±* 20% of their baseline value. This is the range explored in Cone and Shouval (2021) (see their Figure 5), but only qualitative results of the population activities were reported and only for a subset of all possible combinations. In the baseline network, on average 17 out of 50 reported recall times were off by *±* 140 ms (or 20% of correct interval) when measured relative to their expected onset time, whereas these values varied between 15 and 22 for the tested parameter configurations (see Figure 3A, top left). Averaged across all four columns, the outliers decreased to a range between 11-15 (Figure 3A, bottom left). Next, we used a modified z-score based on the median absolute deviation (Iglewicz and Hoaglin, 1993) to evaluate the distribution of the absolute recall times (not relative to their expected onset). These were centered closely around the mean recall time in each column, with the number of outliers decreasing significantly to below 1.5 (3% of recall trials, Figure 3A, right). These results suggest that the recall times are relatively consistent for each column (narrowly distributed), but the absolute deviations from the expected values increase with the element’s position in the sequence.

In other words, the errors and variability accumulate with sequence length, with the network being particularly sensitive to the weaker excitatory connections from Timer onto Messenger cells (see ∆*w* = *−*20% for *T→ M*). In fact, these errors manifest in recalling increasingly shorter intervals (Figure 3B, left), with the last column reporting on average close to 600 ms instead of 700 ms. Averaged across all columns, the median recall time is more accurate. Similar results are obtained for variations in the inhibitory projections between columns (Figure 3B, right).The model displays similar robustness to variations in the eligibility trace time constants 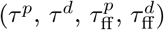 and the variables scaling the Hebbian contribution to the trace dynamics (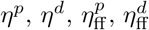, see Methods). Whereas in the original work this analysis was performed with a sequence of two elements of 500 ms each (see Figure 5 - supplement 1 in Cone and Shouval, 2021), here we use a sequence of four 700 ms elements. Compared to the baseline network (Figure 3C, left), where the median recall time decays only slightly with sequence length, randomizing the learning parameters in each learning trial not only increases the median recall time across all columns, but it also leads to a greater variability in the replayed sequences (Figure 3C, center). Randomizing the learning parameters once per network instance does, on average, lead to results closer to the baseline but further increases the recall variability in the last column (Figure 3C, right - analysis not performed in Cone and Shouval, 2021).

These results demonstrate that the system copes well with intermediate perturbations to the baseline parameters with respect to the afferent weights for the Messenger population, the cross-columnar inhibition and the learning rule variables.

While the Timer and Messenger cells are responsible for maintaining a sequence element in the activity and signaling the onset of subsequent ones, the dynamics of the inhibitory populations orchestrates the timing of the individual components. For example, through their characteristic activity curve, the inhibitory cells in *L*_5_ simultaneously control the activity of the Messenger cells in their own column and the onset of the Timer populations in the next column. By modifying the synaptic weight from the Timer cells to the inhibitory population in their column 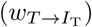, and thus controlling direct excitation, we sought to understand how these inhibitory cells impact learning.

For values significantly lower than baseline (*< −* 25%, grey area in Figure 4A), the network fails to recall sequences in a reliable manner (Figure 4B), in particular sequences containing more than two elements. In addition, the recall times vary significantly across the columns in the case of reduced weights. As the weights increase, the stronger net excitation causes longer-lasting inhibition by *I*_L_5, delaying the activation of the Messenger cells (Figure 4C). This leads to an over-estimation of the elements’ duration, which increases with the element’s position in the sequence (up to +200 ms for 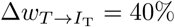, Figure 4D).Although these observations suggest a robust learning mechanism, they also indicate an intrinsic and consistent bias of the model for reporting increasingly shorter intervals and larger variability in the recall times of longer sequences.

### 2.4 Model scaling

In the previous section we investigated the sensitivity of the model to the choice of synaptic weights, but a broader definition of robustness also encompasses invariance to the size of the different populations. Ideally, the model should retain its dynamical and learning properties also for larger network sizes, without the need for manual recalibration of the system parameters. In balanced random networks, increasing the network size by a factor of *m* and decreasing the synaptic weights by a factor of 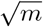 should maintain the activity characteristics (van Vreeswijk and Sompolinsky, 1998; Litwin-Kumar and Doiron, 2012; van Albada et al., 2015). The model studied here differs significantly from these systems with respect to features such as the ratio of excitation and inhibition (1:1, not 4:1), or strong recurrent connectivity in the small *N* regime, which results in significant fluctuations driven by noise. Furthermore, the stereotypical activation patterns underlying sequence learning and replay are significantly more complex. These considerations suggest that successful scaling may require additional modifications of the connectivity.

In the original formulation of the model, each population (Messenger, Timer, inhibitory) consists of 100 neurons. To study how well the model scales for *N*^*′*^ = 400, we kept all parameters unchanged and scaled all non-plastic weights by 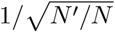 (see Supplementary Table S4). Under such standard scaling, the system fails to learn and recall sequences (Figure 5A), primarily due to the high firing rates of *I*_T_ cells. These decay slower than the corresponding Timer cells, inhibiting the Timer population in the subsequent column and thus prohibiting a correct sequential activation during training.T

Nevertheless, it is possible to find a set of parameters (see Methods and Supplementary Table S4) for which learning unfolds as expected; this is illustrated in Figure 5B. The critical component here is the activity of *I*_T_ (see also Figure 4). This must fulfil three criteria: first, it must decay slightly faster than the rate of the Timer population in the same column; second, it must sufficiently inhibit the Timer populations in all other columns to enable a WTA dynamics; third, the WTA inhibition of the Timer populations must be weak enough that they can still be activated upon stimulation. One way to achieve this is by further decreasing the local weights 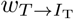 within a column and the cross-columnar inhibition 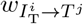. This indicates that, given the right set of parameters, the dynamics underlying the learning process are independent of the network size. Although it is outside the scope of this work, scaling can be likely achieved for a wider range of model sizes, as long as the core properties described above are retained.

### 2.5 Projections between all columns

In the original implementation of Cone and Shouval (2021), and in contrast to the description in the paper, excitatory projections between columns were only allowed in a feedforward manner, thus hard-wiring the order of the sequence elements. Since such a predetermined and stimulus-dependent connection pattern weakens the model’s claims of biological plausibility, we probed the model’s ability to learn when this constraint was relaxed.

To this end, we extended the baseline network with additional projections from Messenger cells in column *C*_*i*_ to Timer cells in all other columns *C*_*j*_, (*i* = *j*) as depicted in Figure 6A. As the weights of these projections are initialized close to 0, no further measures were necessary to maintain the same activity level as the baseline network. Although learning initially proceeded as before, the activity soon lost its stereotypical temporal structure and the learning process is corrupted (Figure 6B). After only a few dozen trials, the activation order of the columns did not match the stimulation, with multiple populations responding simultaneously. Such random, competitive population responses also continued throughout the recall trials.

**Figure 6:**
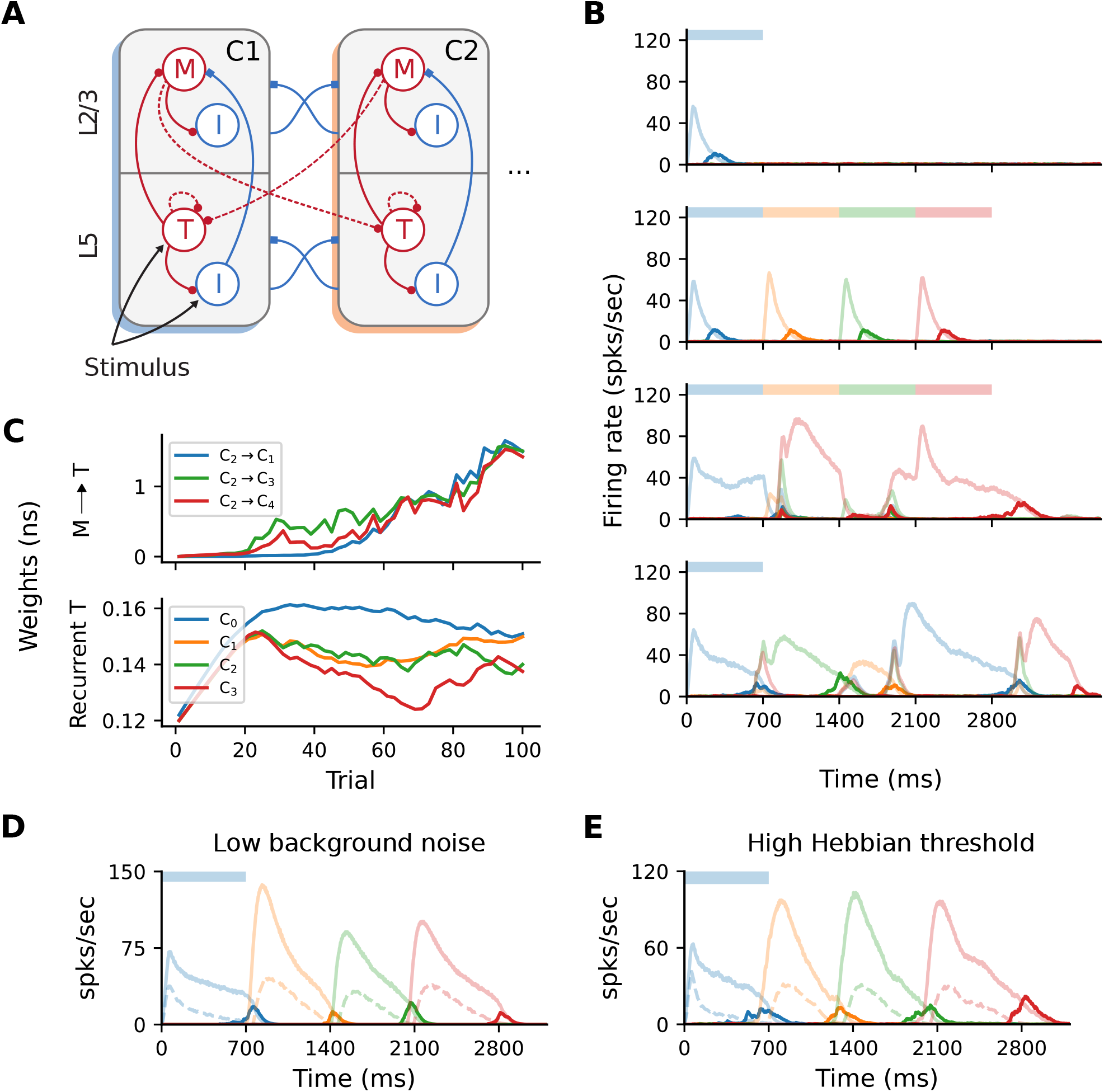
All-to-all cross-columnar excitation prohibits learning. **(A)** Extending the original architecture described in Figure 1B, *M → T* connections exist between all columns *C*_*i*_ *→ C*_*j*_ (*i* = *j*) and are subject to the same plasticity. **(B)** Firing rates of the excitatory populations during learning and recall of four time intervals (each 700 ms). Initially, learning evolves as in Figure 1C, but the activity becomes degenerated and the sequence can not be recalled correctly (lower panels). **(C)** Evolution of the cross-columnar (from *C*_2_, top panel) and recurrent Timer synaptic weights (bottom panel). The transition to the next sequence cannot be uniquely encoded as the weights to all columns are strengthened. **(D)** Sequence recall after 100 training trials in a network with a low background noise (50% of the baseline value, 1*/*2*σ*_*ξ*_). **(E)** Sequence recall after 100 training trials in a network with a higher Hebbian activation threshold for the cross-columnar projections 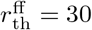 spks*/*sec (instead of the baseline 20 spks*/*sec).

This behavior arises because projections from the Messenger cells to all columns are incorrectly strengthened, not just between subsequent ones reflecting the order of the input sequence. Figure 6C illustrates such an example, with synaptic weights from Messenger cells in *C*_2_ to all other columns *C*_*j*_ being equally strengthened, instead of only to *C*_3_. Naturally, this effect is detrimental because Messenger cells can activate multiple Timer populations at once, introducing a stochasticity in the network that abolishes the unique sequential activation required for accurate learning and recall. In other words, the physical pathway encoding the transitions between sequence elements can not be uniquely traced out as in the baseline network.

According to the Hebbian-based plasticity rule (see Methods), synaptic weights are modified during the reward period only if there is a co-activation of the pre- and postsynaptic neurons. This means that connections from *M* cells in a column *C*_*i*_ to *T* cells in any *C*_*j*_ may be strengthened if there is temporal co-activation of the two populations. While this is the intended behavior for subsequent columns *C*_*i*_ and *C*_*i*+1_, Timer cells in other columns may also spike due to the background noise, thereby enhancing the corresponding connections. Obviously, in the pre-wired (baseline) network this is not an issue, as only subsequent columns are connected.

One straightforward solution to overcome this problem is to reduce the background noise below the spiking threshold, thereby ensuring that only the stimulated populations are active and no “cross-talk” occurs through spurious spiking. Doing so allows the network to regain its functional properties (Figure 6D), pending some minor additional parameter tuning (see Methods). However, from the point of view of biological plausibility, this has the disadvantage that neurons spike exclusively during their preferred stimulus.

Alternatively, it is possible to compensate for the low-rate spontaneous spiking by raising the activation threshold for the Hebbian term, 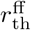 (see Methods). For instance, increasing from the baseline value of 20 to 30 spks*/*secis sufficient to ensure that only the stimulated populations reach these rates. Thus, only synapses between stimulated populations are modified, and the learning process is not affected (Figure 6E). The role and plausibility of such thresholds is detailed in the Discussion.

### 2.6 Alternative wiring with local inhibition

Unlike cortical circuits, where inhibition is assumed to be local (Douglas and Martin, 2004; Fino and Yuste, 2011; Tremblay et al., 2016), the original architecture described in Figure 1B relies on (long-range) inhibitory projections between columns to ensure a soft WTA mechanism in the presence of background activity. This aspect is briefly discussed in Cone and Shouval (2021), and the authors also propose an alternative, biologically more plausible and functionally equivalent network architecture (see their Figure 9). As schematically illustrated in Figure 7A, cross-columnar inhibition can be replaced by local inhibition and corresponding excitatory projections onto these circuits. In contrast to the baseline network, where both Timer and inhibitory cells in *L*_5_ were stimulated, here only Timer cells received input. Otherwise, excitation onto *I*_T_ would soon silence the Timer cells, prohibiting the longer timescales required for encoding the input duration.

**Figure 7:**
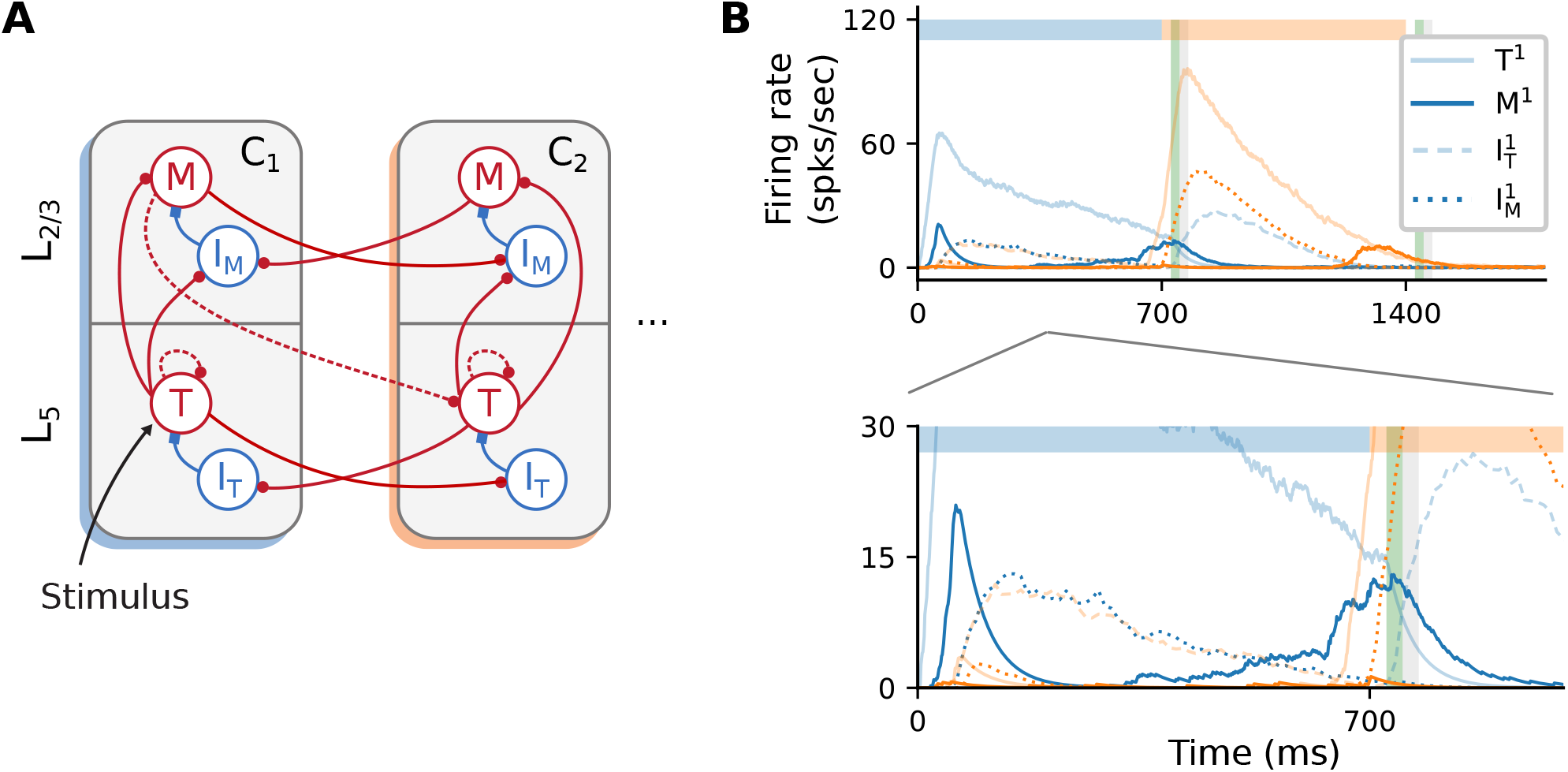
Alternative wiring with local inhibition and only excitatory cross-columnar projections. **(A)** Architecture with local inhibition functionally equivalent to Figure 1B. Inhibitory projections are now local to the column, and feedforward inhibition is achieved via cross-columnar excitatory projections onto the *I* populations. **(B)** Recall of a sequence composed of two 700 ms intervals. Inset (bottom panel) zooms in on the activity at lower rates. As before, color codes for columns. Color shade represents populations in *L*_5_ (light) and *L*_2*/*3_ (dark), with solid curves denoting excitatory populations. Dashed (dotted) curves represent the inhibitory cells *I*_T_ (*I*_M_).

As a proof-of-concept, we empirically derived a set of parameters (see Supplementary Table S5) for such a circuit and found that the core network dynamics and learning process can, in principle, be retained (Figure 7B). However, a significant discrepancy from the baseline behavior concerns the initial transient of the Messenger cells in the first column *C*_1_ (solid, dark blue curve in Figure 7B, bottom panel). This occurs because inhibition onto the Messenger cells from *I*_M_ (dotted, dark blue curves) is slower (due to higher firing threshold) than the excitation from the Timer cells. This results in a brief period of higher Messenger activity before inhibition takes over and silences it. Although this behavior is different from the baseline model, it does not appear to impact learning, and it is in fact consistent with the experimental data from the primary visual cortex (Liu et al., 2015).

## 3 Discussion

Given that the ability to learn and recall temporal sequences may be a universal functional building block of cortical circuits, it is paramount that we understand how such computational capacities can be implemented in the neural substrate. While there have been numerous approaches to model sequence processing in spiking networks, many of these are either unable to capture important functional aspects (e.g., order and duration of sequences), or rely on biophysically unrealistic assumptions in their structure or learning rules. In this work we investigated a recent model proposed by Cone and Shouval (2021), which attempts to overcome these weaknesses. Since here we focused particularly on the reproducibility and replicability aspects, our work provides only limited improvements over the original model. Thus, major modifications such as changes to the learning rule or the evaluation of more complex sequence learning tasks are beyond the scope of our study. However, by re-implementing the model in the NEST simulator, we were able to qualitatively replicate the main findings of the original work, find some of the critical components and assumptions of the model, and highlight its strengths and limitations. More importantly, we provide a complete set of parameters and implementation details for a full replication of the model. As computational studies are becoming increasingly significant across many scientific disciplines, ease of reproduction and replication becomes an ever more important factor, not just to allow efficient scientific progress, but also to ensure a high quality of the work. These points are well illustrated by a notable outcome of this study: as a result of our findings, the authors of the the original study have modified their published code to enable full replication and correct the inconsistencies and errors discovered in their work, as listed below.

### 3.1 Reproducibility

The original model is described in Cone and Shouval (2021), with most parameters provided as Supplementary Information, along with a publicly available MATLAB implementation on ModelDB ^1^. However, while the results are reproducible using the provided implementation in the R^3^ sense described by Benureau and Rougier (2018), a successful replication in the R^5^ sense would not have been possible based solely on the information in the manuscript and Supplementary Tables, given that a number of parameters are either under-specified or omitted entirely. Table 1 and Table 2 give an overview of the more important discrepancies between the description and original implementation, categorized by the their relevance and type of mismatch.

**Table 1:**
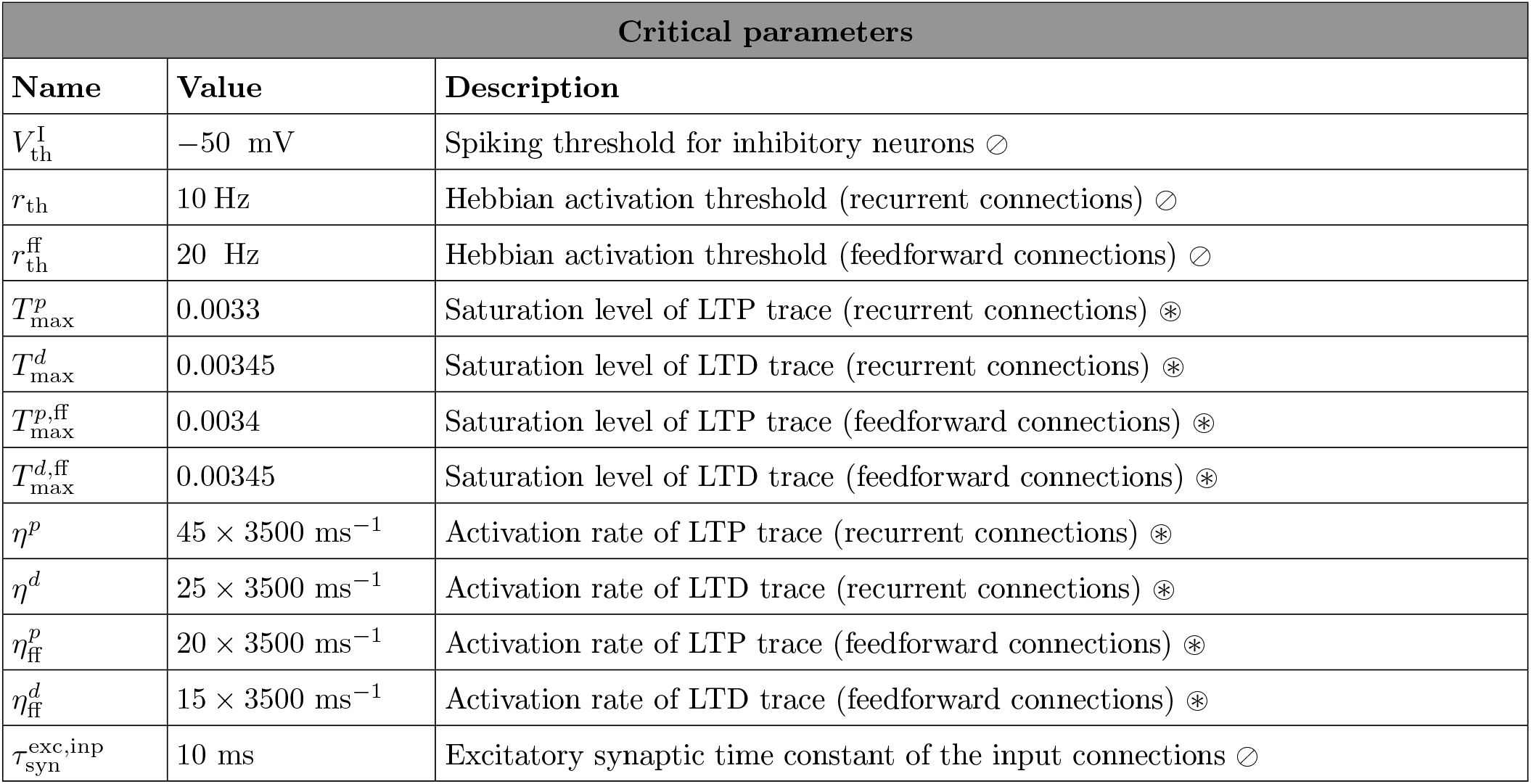
Critical parameters necessary for accurate learning. Symbols denote different discrepancy types: ⊘ represents parameters not mentioned in the study, and ⊛ parameters with only relative but no exact valuesgiven

| Critical parameters |  |  |
| --- | --- | --- |
| Name | Value | Description |
| $V_{\text{th}}^{\text{I}}$ | $-50 \text{ mV}$ | Spiking threshold for inhibitory neurons $\odot$ |
| $r_{\text{th}}$ | 10 Hz | Hebbian activation threshold (recurrent connections) $\odot$ |
| $r_{\text{th}}^{\text{ff}}$ | 20 Hz | Hebbian activation threshold (feedforward connections) $\odot$ |
| $T_{\text{max}}^p$ | 0.0033 | Saturation level of LTP trace (recurrent connections) $\otimes$ |
| $T_{\text{max}}^d$ | 0.00345 | Saturation level of LTD trace (recurrent connections) $\otimes$ |
| $T_{\text{max}}^{p,\text{ff}}$ | 0.0034 | Saturation level of LTP trace (feedforward connections) $\otimes$ |
| $T_{\text{max}}^{d,\text{ff}}$ | 0.00345 | Saturation level of LTD trace (feedforward connections) $\otimes$ |
| $\eta^p$ | $45 \times 3500 \text{ ms}^{-1}$ | Activation rate of LTP trace (recurrent connections) $\otimes$ |
| $\eta^d$ | $25 \times 3500 \text{ ms}^{-1}$ | Activation rate of LTD trace (recurrent connections) $\otimes$ |
| $\eta_{\text{ff}}^p$ | $20 \times 3500 \text{ ms}^{-1}$ | Activation rate of LTP trace (feedforward connections) $\otimes$ |
| $\eta_{\text{ff}}^d$ | $15 \times 3500 \text{ ms}^{-1}$ | Activation rate of LTD trace (feedforward connections) $\otimes$ |
| $\tau_{\text{syn}}^{\text{exc},\text{inp}}$ | 10 ms | Excitatory synaptic time constant of the input connections $\odot$ |

**Table 2:**
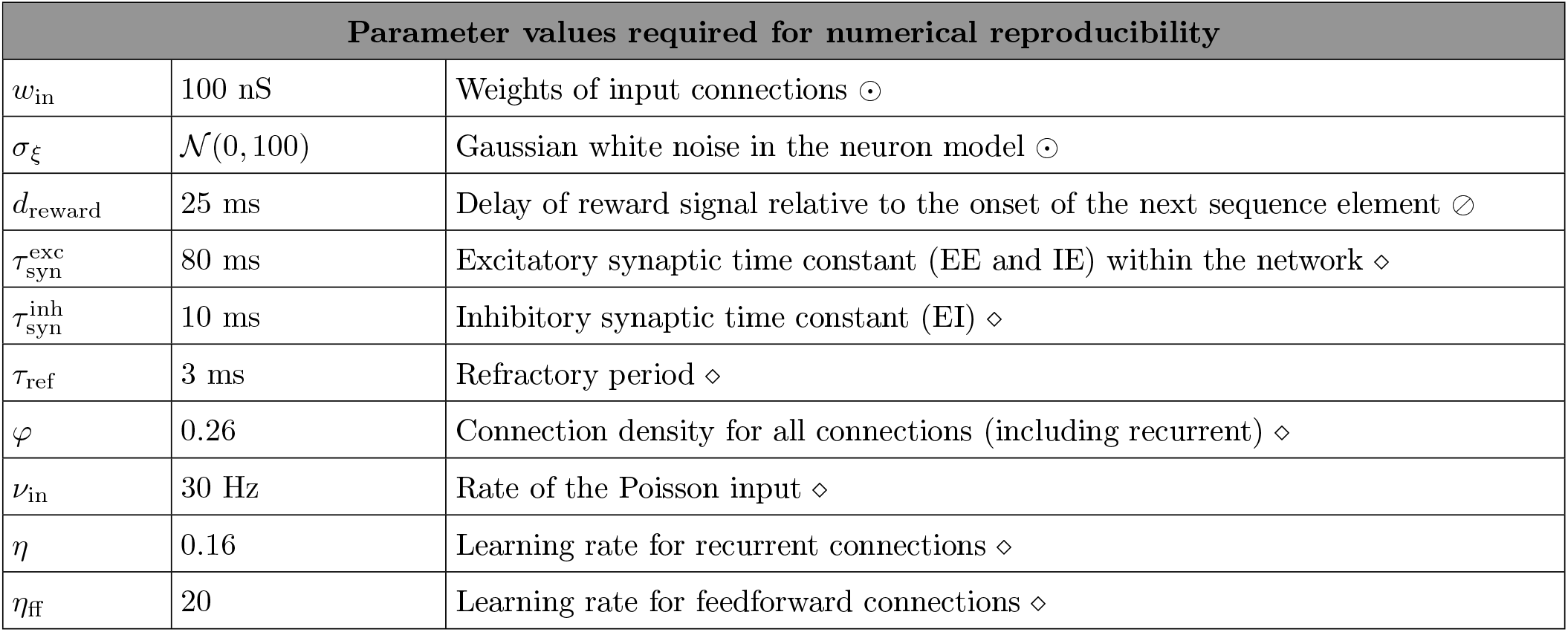
Parameter values needed for obtaining numerically similar results to those reported in Cone and Shouval (2021). Symbols ⊘ and ⊛ as in Table 1. Additionally, ⊙ denotes parameters with no specific values given, while ⋄ denotes a mismatch between the values reported in the paper and the ones used in the reference implementation.

Table 1 lists omitted (or inaccurately stated) critical parameters, i.e. those that are necessary for the model to carry out the computational tasks that are central to the original study. Such oversights are particularly problematic, as they not only make replication more challenging, but also make implicit model assumptions opaque. An illustrative example of an omitted critical parameter is the spiking threshold for the inhibitory neurons, *V*_th_, which is 5 mV higher than the threshold for the excitatory neurons. This is important, as it results in the inhibitory rates decaying slightly faster than the Timer cells, thus activating the Messenger cells at the appropriate time. In the absence of this dynamical feature, learning fails (see for example Figure 5A). While there is some experimental evidence for such a difference in the spiking threshold, it varies significantly across different cell types and recording locations (Tripathy et al., 2015). Similarly, the activation thresholds for the Hebbian learning, *r*_th_, are necessary to ensure that spontaneous spiking resulting from the neuronal noise does not lead to potentiation of unwanted synapses, in particular if connections between all columns are allowed (see Figure 6). Without such thresholds, learning still converges in the baseline network, but the fixed point of the feedforward weights is shifted, stabilizing at a lower value than in the baseline system (see Supplementary Figure S2). Therefore, the role and optimal value for the thresholds likely depends on the amount of noise and spontaneous activity in the network.

A further example is the parameterization of the eligibility traces. Whereas the time constants of the eligibility traces determine their rise and decay behavior, the saturation levels 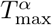 can profoundly impact learning. Forthe Timer cells, although their exact values (not provided in the original work) is not essential, the order of magnitude is still critical; they must be carefully chosen to ensure that the traces saturate soon after stimulus onset, and the falling phase begins before the next reward period (see also Huertas et al., 2015). In other words, even though the parameter space is underconstrained and multiple values can lead to accurate learning Huertas et al. (2016), these nevertheless lie within a restricted interval which is difficult to determine given only the relative values as in the original work: for instance, a value of 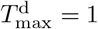 and 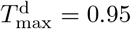 will lead to an abrupt increase in the recurrent Timer weights and learning fails. If the traces do not saturate, learning becomes more sensitive to the trace time constants and the range of time intervals that can be learned with one set of parameters shrinks significantly. Moreover, the excitatory input synapses have a shorter time constant of 10 ms than in the rest of the network, which is required for the fast initial ramp-up phase of the Timer cell activity.

Table 2 summarizes other, less critical parameters, which are nonetheless necessary to achieve qualitatively similar activity levels to those presented in the original work. These include input related parameters (input weights, input rate), as well as the neuronal noise. Whereas some of these discrepancies are due to omission (e.g., noise) or mismatch between the reported and used values (e.g., learning rate), others arise from tool- and implementation particularities. For instance, for *N* = 100 the random number generation in MATLAB results in an effective connectivity *φ* =*∼* 0.26 instead of the 0.3 reported in Cone and Shouval (2021), while the effective refractory period is 3 instead of 2 ms, as threshold crossings are registered with a delay of one simulation step. Although these parameters influence the level of the activity in the network, they do not directly impact the learning process; the key computational features claimed for the model are maintained.

### 3.2 Learning cross-columnar projections

One of the key properties of the model is the ability to learn the order of temporal sequences, achieved by learning the transitions between stimulus-specific populations encoding the sequence elements. However, Cone and Shouval (2021) state that “Messenger cells can only learn to connect to (any) Timer cells outside of their column”, which we interpret as an assertion that Timer cells make connections to Messenger cells in all other columns. In practice, the authors’ reference implementation restricts these to subsequent columns only. This means that the order of the sequence is hardwired into the connectivity, and the system is only learning the duration of the elements. As we demonstrated in Section 2.5, with the baseline parameters the network fails to learn if this restriction is relaxed and feedforward projections are indeed allowed between any columns.

A simple way to circumvent this problem is to ensure that neurons outside the populations coding for the current stimulus remain completely (or sufficiently) silent, so as to avoid the co-activation necessary for Hebbian synaptic potentiation (see Figure 6D). Although such an idealized behavior may be an appropriate solution from a modelling perspective, neurons in the cortex are rarely tuned exclusively to particular stimuli. Instead, most cells spike irregularly (typically at a low rate) even in the absence of input (ongoing activity, see e.g., Arieli et al., 1996), and many respond to multiple different inputs (Walker et al., 2011; Rigotti et al., 2013; de Vries et al., 2020).

A biologically more plausible alternative is to increase the Hebbian activation threshold *r*_th_, such that noise-induced spontaneous activity does not lead to a modification of the synaptic strength. However, this introduces an additional, critical parameter in the model. Furthermore, such hard thresholds are coupled to the intensity of background activity and spontaneous spiking, with occasional higher rates possibly destabilizing the learning process.

### 3.3 Functional and neurophysiological considerations

From a functional perspective, a generic model of sequence processing should be able to perform various related tasks in addition to sequence replay, such as chunking, learning compositional sequences and handling non-adjacent dependencies in the input (Fitch and Martins, 2014; Wilson et al., 2018; Hupkes et al., 2019). Although Cone and Shouval (2021) discuss and provide an extension of the baseline network for higher-order Markovian sequences, the computational capacity of the model is fundamentally limited by the requirement of a unique stimulus-column (or stimulus-population) mapping. This characteristic means that for certain tasks, such as learning (hierarchical) compositional sequences (i.e., sequences of sequences), the model size would increase prohibitively with the number of sequences, as one would require a dedicated column associated with each possible sequence combination. In addition, it would be interesting to evaluate the model’s ability to recognize and distinguish statistical regularities in the input in tasks such as chunking, which involve one or more sequences interleaved with random elements.

In their study, Cone and Shouval (2021) demonstrate that the extended, rate-based network can learn multiple, higher-order Markovian sequences when these are presented successively. For first-order Markovian sequences, this should also hold for the baseline spiking network model, contingent on preserving the unique stimulus-to-column mapping. However, it is also important to understand how the model behaves when two sequences are presented *simultaneously*. This depends on the interpretation and expected behavior, and to the best of our knowledge there is little experimental and modeling work on this (but see, e.g., Murray and Escola, 2017). Nevertheless, if the two sequences are considered to be *independent*, we speculate that the networks will not be able to learn and treat them as such for multiple reasons. Assuming that projections between all columns are allowed (with the appropriate measures, see Section 2.5), in the spiking model the connections between the columns associated with the different sequences would also be strengthened upon temporal co-activation: for two simultaneously initiated sequences S1 and S2, the cross-columnar projections between a column 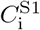 associated with S1 and another column 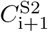coding for an element at position *i* + 1 in S2 would be (incorrectly) strengthened. In the case of the extended rate network, the context representations may mix and interfere in the external reservoir, and the issue of temporal co-activation discussed above is also likely to occur.

Moreover, convergence of learning in the cross-columnar synapses depends on the existence of two consecutive reward periods. As described in Section 2.2 and illustrated in Figure 2C, during the first reward (associated with the current sequence element) the weights are potentiated, even after the weights have reached a fixed point. However, a second reward, during which the weights are depressed, is necessary to achieve a net zero difference in the LTP and LTD traces at lower weight values. Although learning would converge even without a second reward, the fixed point will be different (higher), and thus convergence would occur for larger weights (possibly too large for stable firing rates). Given that the reward (novelty) signal is globally released both before and after each sequence element in the interpretation of Cone and Shouval (2021), the existence of a reward after the final element is guaranteed and therefore this is not an issue for the stimulation protocol used in the original and our study. If, on the other hand, we interpret the reward as a novelty signal indicating the next stimulus, we would not expect it to be present in this form after the last element of the sequence. In this case, the cross-columnar projections marking the transition from the penultimate to the ultimate element may not be learned accurately (weights would still converge, but likely to larger values than appropriate).

While a solution to the above issues is beyond the scope of this work, we speculate that a more granular architecture, in which multiple stimulus-specific sub-populations could form different cell assemblies within a single column, would be more in line with experimental evidence from the neocortex. Some functional specialization of single cortical columns has been hypothesized (Mountcastle, 1997; Harris and Shepherd, 2015), but such columns are typically composed of a number of cell groups responsive to a wider range of stimuli. We assume that mapping the model to such an extended columnar architecture would require a more complex, spatially-dependent connectivity to ensure similar WTA dynamics. The requirement of completely segregated populations tuned to unique stimuli, however, is more difficult to overcome and reconcile with experimental data. While the tuning curves of many cells (but by far not all, see de Vries et al., 2020) in the early sensory cortices are indeed strong and sharp (Hubel and Wiesel, 1959; Bitterman et al., 2008), these become weaker and broader in the following stages of the cortical hierarchy, where cells typically exhibit mixed selectivity (Rigotti et al., 2013; Fusi et al., 2016). Thus, more complex tasks requiring a mixture of representations can not be easily conceptualized in the context of the proposed network architecture.

As we demonstrated in Section 2.6, the model is relatively flexible with respect to the precise wiring patterns, as long as certain core, inhibition-related properties are preserved. Given that long-range projections in the neocortex are typically excitatory (Brown and Hestrin, 2009; Douglas and Martin, 2004), the original architecture (see Figure 1B) was implausible due to its reliance on cross-columnar inhibition. The relative ease in adapting the wiring to have only local inhibition is indicative of simple yet powerful and modular computational mechanisms, suggesting that these may be used as building blocks in more complex sequence learning architectures.

Despite these limitations and sensitivity to some parameters, the model presented by Cone and Shouval (2021) is an important step towards a better understanding of how cortical circuits process temporal information. While its modular structure enabling spatially segregated representations may be more characteristic for earlier sensory regions, the proposed local learning rule based on rewards, partially solving the credit assignment problem, is a more universal mechanism likely to occur across the cortex.

## 4 Materials and methods

The sequence learning model analyzed in this study is described in full detail in the original work of Cone and Shouval (2021). Nevertheless, given the numerous discrepancies between the model description and implementation (see Discussion), we present all the key properties and parameters that are necessary for a successful replication of the results, including the extended architectures investigated in Section 2.5 and Section 2.6.

### 4.1 Network architecture

The central characteristic of the network architecture is the modular columnar structure (see Figure 1A, B), where each of the *N*_C_ columns is associated with a unique sequence element (stimulus). Each column contains two excitatory (Timer and Messenger) and two associated inhibitory populations *I*_T_and *I*_M_, roughly corresponding to *L*_5_ and *L*_2*/*3_ in the cortex. In the following, we will refer to these cell populations as 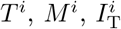 and 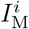,respectively, where the superscript *i* denotes the column *C*_*i*_.

Each of the above populations is composed of *N* = 100 leaky integrate-and-fire neurons, with the exception of the network simulated in Section 2.4, where *N* = 400. The wiring diagram of the baseline network used in Cone and Shouval (2021) is schematically illustrated in Figure 1B. Within a column *C*_*i*_, *T*^*i*^ cells connect to 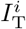and *M*^*i*^, in addition to recurrent connections to other *T*^*i*^ cells. *M*^*i*^ neurons excite the local inhibitory population 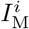, and are inhibited by 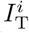. Inhibition onto the excitatory cells also exists between the columns in a layer-specific manner i.e., 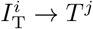 and 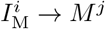, with *i* ≠ *j*. Lastly, *M*^*i*^ cells in *C*_*i*_ connect in a feedforward manner to *T*^*i*+1^ cells in the subsequent column *C*_*i*+1_. All connections within the same and between different populations have a density of *φ* = 0.26. Note that only the feedforward projections *M*^*i*^ *→T*^*i*+1^ and the recurrent *T*^*i*^*→ T*^*i*^ connections are subject to plasticity (see below); all other connections are static. The plastic weights are initialized close to 0 and the static weights are normally distributed around their mean values with a standard deviation of 1.The complete set of parameters for the architecture proposed in Cone and Shouval (2021) as well as the variants described below are specified in the Supplementary Materials.

#### 4.1.1 Scaled model

For the scaled network model described in Section 2.4, the number of neurons in each populations was increased to *N*^*′*^ = 400 from *N* = 100. To keep the input variance constant, in the standard scaling scenario (Figure 5A) we followed the common approach for balanced random networks (van Vreeswijk and Sompolinsky, 1998; Litwin-Kumar and Doiron, 2012) and reduced all non-plastic synaptic weights by multiplying them with 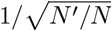. In addition, we halved the standard deviation *σ*_*ξ*_ of the background noise such that the firing rates were in the same range as for the baseline network. To restore the functional aspects of the network, additional tuning was required for most of the projections, see Supplementary Table S4.

#### 4.1.2 All-to-all cross-columnar connectivity

In Section 2.5, the baseline network is modified by instantiating plastic excitatory connections between all columns *M*^*i*^ *→ T*^*j*^, (*i* ≠ *j*) rather than solely between the columns representing consecutive elements of the stimuli (see Figure 6A). All other parameters are unchanged.

#### 4.1.3 Alternative wiring with local inhibition

The functionally equivalent network analyzed in Section 2.6 required multiple modifications (see Figure 7A). Inhibitory connections are local to the corresponding layer, with connections 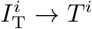 and 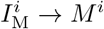. Timer cells *T*^*i*^ project to both *M*^*i*^ and 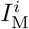, as well as to 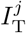 in other columns *C*_*j*_. In layer *L*_2*/*3_, *M*^*i*^ cells project to*T*^*i*+1^ and 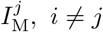.

### 4.2 Neuron model

The networks are composed of leaky integrate-and-fire (LIF) neurons, with fixed voltage threshold and conductance-based synapses. The dynamics of the membrane potential *V*_*i*_ for neuron *i* follows:

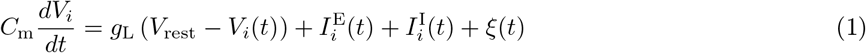

where the leak-conductance is given by *g*_L_, 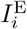 and 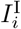 represent the total excitatory and inhibitory synaptic input currents, and *ξ* is a noise term modelled as Gaussian white noise with standard deviation *σ*_*ξ*_ = 100, unless otherwise stated. This noise term is sufficient to cause a low baseline activity of around 1 − 2 spks*/*sec. Upon reaching a threshold *V*_th_ = *−*55 mV (*−* 50 mV for inhibitory neurons), the voltage is reset to *V*_reset_ for a refractory period of *t*_ref_ = 3 ms. Note that the higher threshold for inhibitory neurons is critical for the faster decay of their activity compared to Timer cells.The dynamics of the synaptic conductances are modelled as exponential functions with an adaptation term, with fixed and equal conduction delays for all synapse types. The equations of the model dynamics, along with the numerical values for all parameters are summarized in Supplementary Tables S1-3.

In all figures depicting firing rates, these are estimated from the spike trains using an exponential filter with time constant *τ*_r_ = 40 ms.

### 4.3 Eligibility-based learning rule

The main assumption of the learning rule is the availability of two synaptic eligibility traces at every synapse 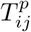and 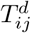, representing long-term potentiation (LTP) and depression (LTD), which can be simultaneously activated through the Hebbian firing patterns.

For *a ∈ {p, d}*, the dynamics of the traces follows:

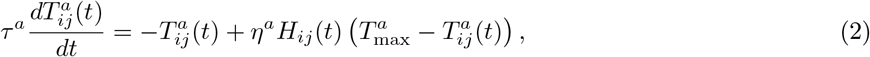

where *τ*^*a*^ is the time constant, *η*^*a*^ is a scaling factor, and 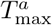 is the saturation level of the trace. *H*_*ij*_(*t*) is the Hebbian term defined as the product of firing rates of the pre- and postsynaptic neurons:

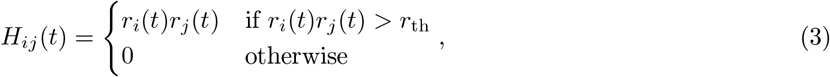

with *r*_th_ 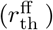 representing different threshold values for recurrent *T* to *T* (feedforward *M* to *T*) connections.Note that while this equation is used in both the original MATLAB implementation and in our re-implementation in NEST, the Hebbian terms in the equations in Cone and Shouval (2021) are further normalized by 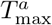. For adetailed analysis of the learning convergence, see the original study.

These activity-generated eligibility traces are silent and transient synaptic tags that can be converted into long-term changes in synaptic strength by a third factor, *R*(*t*) which is modelled here as a global signal using a delta function, *R*(*t*) = *δ*(*t− t*_reward_*− d*_reward_), and is assumed to be released at each stimulus onset/offset. Although typically signals of this sort are used to encode a *reward*, they can also, as is the case here, be framed as a *novelty* signal indicating a new stimulus. Hence, the synaptic weights *w*_*ij*_ are updated through

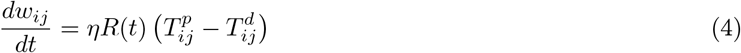

where *η* (*η*_ff_ for feedforward) is the learning rate. Following the reward signal, which has a duration of 25 ms, the eligibility traces are “consumed” and reset to zero, and their activation is set into a short refractory period of 25 ms. In practice, although the weight updates are tracked and evolve during each reward period according to Equation 4, they are only updated at the end of the trial. However, this does not affect the results in any significant manner (data not shown).

### 4.4 Stimulation protocol

Stimulus input is modelled as a 50 ms step signal, encoded as Poisson spike trains with a rate *ν*_in_ = 30 spks*/*sec. In the baseline and the extended network discussed in Section 2.5, this input is injected into both *T*^*i*^ and 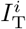 cells, with synaptic weights *w*_in_. In the network discussed in Section 2.6, the input is restricted to *T*^*i*^.

The training process of a network instance consists of 100 trials (unless otherwise stated), and in each trial the corresponding columns are stimulated at certain time points according to the input sequence, with the interval between elements representing the duration of the stimulus. At the beginning of each trial, the state of the neurons (membrane potential) and the eligibility traces are reset to their initial values. The test phase consists of multiple trials (usually 50), where the sequence is replayed upon a cued stimulation of the first column.

### 4.5 Numerical simulations and analysis

All numerical simulations were conducted using the Functional Neural Architectures (FNA) toolkit v0.2.1 (Duarte et al., 2021), a high-level Python framework for creating, simulating and evaluating complex, spiking neural microcircuits in a modular fashion. It builds on the PyNEST interface for NEST (Gewaltig and Diesmann, 2007), which provides the core simulation engine. To ensure the reproduction of all the numerical experiments and figures presented in this study, and abide by the recommendations proposed in (Pauli et al., 2018), we provide a complete code package that implements project-specific functionality within FNA (see Supplementary Materials) using NEST 2.20.0 (Fardet et al., 2020). For consistency checks with the reference implementation, we used *MATLAB* version R2020b.

## Author Contributions

BZ, RD, and AM designed the study. BZ re-implemented the model and performed all simulations and analyses. BZ, RD, and AM contributed to writing of manuscript.

## Acknowledgments

The authors gratefully acknowledge the computing time granted by the JARA-HPC Vergabegremium on the supercomputer JURECA (Jülich Supercomputing Centre, 2021) at Forschungszentrum Jülich. In addition, we would like to thank Charl Linssen of the Simulation and Data Laboratory Neuroscience for his support concerning the neuron and synapse model implementations.

## Supplemental Data

See enclosed Supplementary Material.

## Data Availability Statement

All the relevant data and code is available in a public GitHub repository at https://github.com/zbarni/re_modular_seqlearn. (see also Supplementary Materials). The MATLAB code used in Cone and Shouval (2021) and the revised version can be found in a ModelDB repository at http://modeldb.yale.edu/266774.

## Footnotes

1 http://modeldb.yale.edu/266774

