## Supplementary Material for "Towards reproducible models of sequence learning: replication and analysis of a modular spiking network with reward-based learning"

### 1 Supplementary Figures

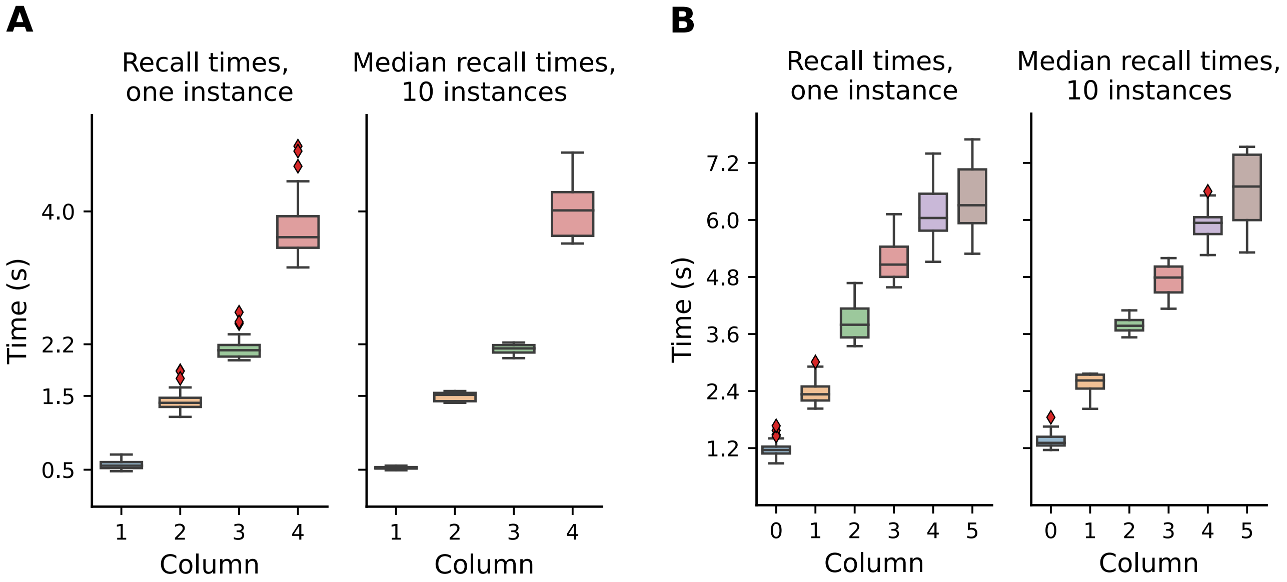


Figure 1: **Fluctuations in learning and recall increase with sequence complexity and number of elements.** **(A)** The network was trained on a sequence of four elements: 500, 1000, 700, 1800 ms. Left: recall times for 30 trials after learning, for one network instance. Right: distribution of the median recall times over 10 network instances, with the median in each network calculated over 30 replay trials. **(B)** Same as (A), for a sequence of six elements with a duration of 1200 ms each.

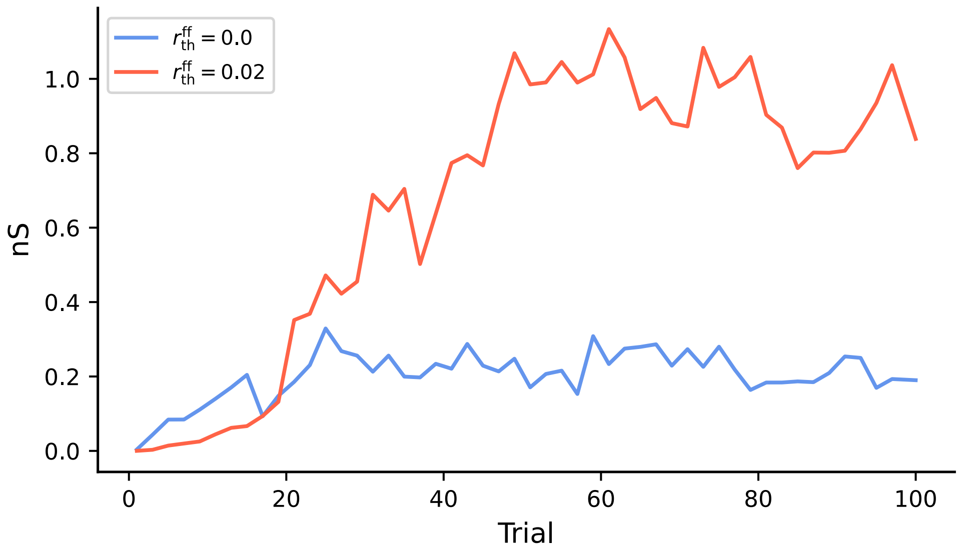


Figure 2: **Hebbian threshold impacts learning convergence of cross-columnar connections.** In the baseline network, learning succeeds even in the absence of a Hebbian threshold $r_{\mathrm{th}}^{\mathrm{ff}}$. While a non-zero threshold leads to larger synaptic weights after convergence, it also increases the variability between trials.

### 2 Supplementary Tables

#### 2.1 Baseline Model

| A: Model Summary | |
|---|---|
| **Populations** | Multiple columns, each one composed of an excitatory Timer (layer $L_5$) and Messenger ($L_{2/3}$) population, with one inhibitory population in each layer |
| **Topology** | None |
| **Connectivity** | Sparse, random recurrent connectivity |
| **Neuron Model** | Leaky integrate-and-fire, fixed voltage threshold, fixed absolute refractory time, no adaptation |
| **Synapse Model** | Conductance-based, exponentially decaying PSCs, static and plastic synaptic weights, fixed delays |
| **Plasticity** | Reward-based plasticity, short-term adaptation |
| **Input** | Stochastic background current and inhomogeneous Poisson spikes onto stimulus-specific $T$ and $I_{\mathrm{T}}$ |
| **Measurements** | Spiking activity |

| B: Populations | | |
|---|---|---|
| **Name** | **Elements** | **Size** |
| $T^i, I_{\mathrm{T}}^i, M^i, I_{\mathrm{M}}^i$ in column $C_i$ | LIF neuron | 100 |

| C: Neuron Models | |
|---|---|
| **Name** | Leaky integrate-and-fire (LIF) neuron |
| **Subthreshold Dynamics** | if $(t > t^* + \tau_{\mathrm{ref}})$<br>$C_{\mathrm{m}} \frac{dV_i}{dt} = g_{\mathrm{L}} \left( V_{\mathrm{rest}} - V_i(t) \right) + I_i^{\mathrm{E}}(t) + I_i^{\mathrm{I}}(t) + \xi(t)$<br>else<br>$V(t) = V_{\mathrm{reset}}$<br>$I_{\mathrm{ij}}^{\mathrm{syn}}(t) = g_{\mathrm{ij}}^{\mathrm{syn}}(E_{\mathrm{syn}} - V_i(t))$ |
| **Spiking** | If $V(t-) < V_{\mathrm{th}}$ OR $V(t+) \geq V_{\mathrm{th}}$<br>1. set $t^* = t$ 2. emit spike with time stamp $t^*$ |

| D: Synapse Models | |
|---|---|
| **Synaptic trace** | $\frac{\mathrm{ds_i}}{dt} = -\frac{\mathrm{s_i}}{\tau_{\mathrm{s}}} + \rho \left(1 - \mathrm{s_i}\right) \sum_{\mathrm{k}} \delta \left(\mathrm{t} - \mathrm{t_k^i}\right)$ |
| **Name** | Reward-based, with separate LTP and LTD eligibility traces |
| **Trace update rule** | $\tau^a \frac{dT_{ij}^a(t)}{dt} = -T_{ij}^a(t) + \eta_{(\mathrm{ff})}^a H_{ij}(t) \left( T_{\max}^a - T_{ij}^a(t) \right), a \in \{p, d\}$<br>$H_{ij}(t) = \begin{cases} r_i(t) r_j(t) & \text{if } r_i(t) r_j(t) > r_{\mathrm{th}}^{(\mathrm{ff})} \\ 0 & \text{otherwise} \end{cases}$ |
| **Online update rule** | $\frac{dw_{ij}}{dt} = \eta_{(\mathrm{ff})} R(t) \left( T_{ij}^p - T_{ij}^d \right)$<br>$R(t) = \delta(t - t_{\mathrm{reward}} - d_{\mathrm{reward}})$ |

| E: Input | | |
|---|---|---|
| **Type** | **Target** | **Description** |
| poisson_generator | $T^i$ and $I_{\mathrm{T}}^i$ in $C_i$ | Total rate $\nu_{\mathrm{in}}$ for a duration of 50 ms |

| F: Measurements |
|---|
| Spiking activity |

Table S1: Tabular description of network model after Nordlie, Gewaltig, and Plesser (2009).

| A: Populations | | | |
|---|---|---|---|
| **Name** | **Value** | **Source** | **Description** |
| $N$ | 100 | paper | Population size of every population, excitatory and inhibitory |

| B: Connectivity | | | |
|---|---|---|---|
| **Name** | **Value** | **Source** | **Description** |
| $d$ | 1 ms | code | Synaptic transmission delay |
| $\varphi$ | 0.26 | code* | Connection probability for all populations |
| $w_{\text{in}}$ | 100 nS | code | Synaptic strength of input connections |
| $w_{T\to M}$ | 0.2 nS | code* | Intracolumnar $T$ to $M$ excitatory synaptic strength |
| $w_{I_\text{T}\to M}$ | 70 nS ⊛ | code* | Inhibitory synaptic strength from $I_\text{T}$ in $L_5$ to $M$ |
| $w_{I^i_\text{T}\to T^j}$ | 100 nS ⊛ | code* | Inhibitory synaptic strength from $I^i_\text{T}$ in $C_i$ to $T^j$ in $C_j$ |
| $w_{I^i_\text{M}\to M^j}$ | 100 nS ⊛ | code* | Inhibitory synaptic strength from $I^i_\text{M}$ in $C_i$ to $M^j$ in $C_j$ |
| $w_{T\to I_\text{T}}$ | 0.2 nS ⊛ | code* | Intracolumnar $T$ to $I_\text{T}$ excitatory synaptic strength |
| $w_{M\to I_\text{M}}$ | 1 nS ⊛ | code* | Intracolumnar $M$ to $I_\text{M}$ excitatory synaptic strength |

| B: Neuron Model | | | |
|---|---|---|---|
| **Name** | **Value** | **Source** | **Description** |
| $C_\text{m}$ | 200 pF | paper | Membrane capacitance |
| $\tau_\text{m}$ | 10 ms | paper | Membrane time constant |
| $g_\text{L}$ | 10 nS | paper | Leak conductance |
| $E_L$ | −60 mV | paper | Resting membrane potential |
| $V^\text{E}_\text{th}$ | −55 mV | paper | Spiking threshold for excitatory neurons |
| $V^\text{I}_\text{th}$ | −50 mV | code* | Spiking threshold for inhibitory neurons |
| $V_\text{reset}$ | −60 mV | code* | Reset potential |
| $\tau_\text{ref}$ | 3 ms | code* | Absolute refractory period |
| $\sigma_\xi$ | 100 | code* | Standard deviation of Gaussian white noise |
| $\nu_\text{in}$ | 30 Hz | code* | Rate of Poisson stimulus input |

| C: Synapse Model | | | |
|---|---|---|---|
| **Name** | **Value** | **Source** | **Description** |
| $E_\text{E}$ | 0 mV | paper | Excitatory reversal potential |
| $E_\text{I}$ | −70 mV | paper | Inhibitory reversal potential |
| $\tau^\text{exc,inp}_\text{syn}$ | 10 ms | code* | Excitatory synaptic time constant of the input connections |
| $\tau^\text{exc}_\text{syn}$ | 80 ms | paper | Excitatory synaptic time constant |
| $\tau^\text{inh}_\text{syn}$ | 10 ms | paper | Inhibitory synaptic time constant |
| $\rho$ | 1/7 | paper | Fractional change of synaptic activation |

Table S2: Tabular description of the neuron, synapse and connectivity parameters. Parameters marked with ⊛ were additionally jittered with a randomly drawn value from $\mathcal{N}(0, 0.1)$. Parameters marked with * had different values in the code than reported in the paper.

| A: Learning Parameters | | | |
|---|---|---|---|
| **Name** | **Value** | **Source** | **Description** |
| $\tau^p$ | 2000 ms | paper | LTP eligibility trace time constant (intracolumnar connections) |
| $\tau^d$ | 1000 ms | paper | LTD eligibility trace time constant (intracolumnar connections) |
| $\tau^p_{\mathrm{ff}}$ | 200 ms | paper | LTP eligibility trace time constant (cross-columnar connections) |
| $\tau^d_{\mathrm{ff}}$ | 800 ms | paper | LTD eligibility trace time constant (cross-columnar connections) |
| $T^p_{\mathrm{max}}$ | 0.0033 | code* | Saturation level of LTP trace (intracolumnar connections) |
| $T^d_{\mathrm{max}}$ | 0.00345 | code* | Saturation level of LTD trace (intracolumnar connections) |
| $T^{p,\mathrm{ff}}_{\mathrm{max}}$ | 0.0034 | code* | Saturation level of LTP trace (cross-columnar connections) |
| $T^{d,\mathrm{ff}}_{\mathrm{max}}$ | 0.00345 | code* | Saturation level of LTD trace (cross-columnar connections) |
| $\eta^p$ | $45 \times 3500$ ms$^{-1}$ | code* | Activation rate of LTP trace (intracolumnar connections) |
| $\eta^d$ | $25 \times 3500$ ms$^{-1}$ | code* | Activation rate of LTD trace (intracolumnar connections) |
| $\eta^p_{\mathrm{ff}}$ | $20 \times 3500$ ms$^{-1}$ | code* | Activation rate of LTP trace (cross-columnar connections) |
| $\eta^d_{\mathrm{ff}}$ | $15 \times 3500$ ms$^{-1}$ | code* | Activation rate of LTD trace (cross-columnar connections) |
| $r_{\mathrm{th}}$ | 10 Hz | code* | Hebbian activation threshold (recurrent connections) |
| $r^{\mathrm{ff}}_{\mathrm{th}}$ | 20 Hz | code* | Hebbian activation threshold (feedforward connections) |
| $\eta$ | 0.16 ms$^{-1}$ | code* | Learning rate $T \to T$ connections |
| $\eta$ | 20 ms$^{-1}$ | code* | Learning rate $M \to T$ connections |
| $T_{\mathrm{reward}}$ | 25 ms | paper | Duration of neuromodulator presentation upon change in stimulus |
| $T_{\mathrm{tr}}$ | 25 ms | paper | Duration of refractory period for traces following neuromodulator presentation |
| $d_{\mathrm{reward}}$ | 25 ms | paper | Reward delay |

Table S3: Tabular description of learning parameters. Parameters marked with * had different values in the code than reported in the paper.

#### 2.2 Scaled model

| **A: Parameters for standard scaling** | | |
|---|---|---|
| **Name** | **Value** | **Description** |
| $N'$ | 400 | Number of neurons in each population (scaled) |
| $w'_{T\to M}$ | $w_{T\to M}/2$ | Intracolumnar $T$ to $M$ excitatory synaptic strength |
| $w'_{I_\mathrm{T}\to M}$ | $w_{I_\mathrm{T}\to M}/2$ $^{\circledast}$ | Inhibitory synaptic strength from $I_\mathrm{T}$ in $L_5$ to $M$ |
| $w'_{I^i_\mathrm{T}\to T^j}$ | $w_{I^i_\mathrm{T}\to T^j}/2$ $^{\circledast}$ | Inhibitory synaptic strength from $I^i_\mathrm{T}$ in $C_i$ to $T^j$ in $C_j$ |
| $w'_{I^i_\mathrm{M}\to M^j}$ | $w_{I^i_\mathrm{M}\to M^j}/2$ $^{\circledast}$ | Inhibitory synaptic strength from $I^i_\mathrm{M}$ in $C_i$ to $M^j$ in $C_j$ |
| $w'_{T\to I_\mathrm{T}}$ | $w_{T\to I_\mathrm{T}}/2$ $^{\circledast}$ | Intracolumnar $T$ to $I_\mathrm{T}$ excitatory synaptic strength |
| $w'_{M\to I_\mathrm{M}}$ | $w_{M\to I_\mathrm{M}}/2$ $^{\circledast}$ | Intracolumnar $M$ to $I_\mathrm{M}$ excitatory synaptic strength |

| **B: Parameters for manually tuned scaling** | | |
|---|---|---|
| **Name** | **Value** | **Description** |
| $N''$ | 400 | Number of neurons in each population (scaled) |
| $w''_{T\to M}$ | $w'_{T\to M}\cdot 1.2$ | Intracolumnar $T$ to $M$ excitatory synaptic strength |
| $w''_{I_\mathrm{T}\to M}$ | $w'_{I_\mathrm{T}\to M}\cdot 2$ $^{\circledast}$ | Inhibitory synaptic strength from $I_\mathrm{T}$ in $L_5$ to $M$ |
| $w''_{I^i_\mathrm{T}\to T^j}$ | $w'_{I^i_\mathrm{T}\to T^j}\cdot 0.02$ $^{\circledast}$ | Inhibitory synaptic strength from $I^i_\mathrm{T}$ in $C_i$ to $T^j$ in $C_j$ |
| $w''_{I^i_\mathrm{M}\to M^j}$ | $w'_{I^i_\mathrm{M}\to M^j}\cdot 2$ $^{\circledast}$ | Inhibitory synaptic strength from $I^i_\mathrm{M}$ in $C_i$ to $M^j$ in $C_j$ |
| $w''_{T\to I_\mathrm{T}}$ | $w'_{T\to I_\mathrm{T}}\cdot 0.02$ $^{\circledast}$ | Intracolumnar $T$ to $I_\mathrm{T}$ excitatory synaptic strength |
| $w''_{M\to I_\mathrm{M}}$ | $w'_{M\to I_\mathrm{M}}\cdot 2$ $^{\circledast}$ | Intracolumnar $M$ to $I_\mathrm{M}$ excitatory synaptic strength |
| $\sigma''_\xi$ | $\sigma_\xi/2$ | Standard deviation of Gaussian white noise |

Table S4: Tabular description of the modified parameters in the scaled network models. For the standard scaling, the values are obtained by applying a scaling factor of $1/\sqrt{N'/N}$ to the original values (see Methods). Parameters marked with $^{\circledast}$ were additionally jittered with a randomly drawn value from $\mathcal{N}(0, 0.1)$.

#### 2.3 Alternative model with local inhibition

| **A: Parameters for Network with Local Inhibition** | | |
|---|---|---|
| **Name** | **Value** | **Description** |
| $N$ | 100 | Number of neurons in each population (as in baseline model) |
| $w_{T \to M}$ | 0.2 nS | Intracolumnar $T$ to $M$ excitatory synaptic strength |
| $w_{I_{\mathrm{T}} \to T}$ | 70 nS$^{\circledast}$ | Inhibitory synaptic strength from $I_{\mathrm{T}}$ in $L_5$ to $T$ |
| $w_{I_{\mathrm{M}} \to M}$ | 70 nS$^{\circledast}$ | Inhibitory synaptic strength from $I_{\mathrm{M}}$ in $L_{2/3}$ to $M$ |
| $w_{T \to I_{\mathrm{M}}}$ | 0.2 nS$^{\circledast}$ | Excitatory synaptic strength from $T$ to $I_{\mathrm{M}}$ |
| $w_{T^i \to I_{\mathrm{T}}^j}$ | 0.2 nS$^{\circledast}$ | Excitatory synaptic strength from $T$ in column $C_i$ to $I_{\mathrm{T}}$ in $C_j, i \neq j$ |
| $w_{M^i \to I_{\mathrm{M}}^j}$ | 0.5 nS$^{\circledast}$ | Excitatory synaptic strength from $M$ in column $C_i$ to $I_{\mathrm{M}}$ in $C_j, i \neq j$ |

Table S5: Tabular description of the modified parameters in the model with rewired local inhibition. Parameters marked with $^{\circledast}$ were additionally jittered with a randomly drawn value from $\mathcal{N}(0, 0.1)$.
